# Clonally distinct differentiation trajectories shape CD8^+^ memory T cell heterogeneity after acute viral infections in humans

**DOI:** 10.1101/832899

**Authors:** Jeff E. Mold, Laurent Modolo, Joanna Hård, Margherita Zamboni, Anton J.M. Larsson, Carl-Johan Eriksson, Patrik L. Ståhl, Erik Borgström, Simone Picelli, Björn Reinius, Rickard Sandberg, Pedro Réu, Carlos Talavera-Lopez, Björn Andersson, Kim Blom, Johan K. Sandberg, Franck Picard, Jakob Michaëlsson, Jonas Frisén

## Abstract

CD8^+^ T cells play essential roles in immunity to viral and bacterial infections, and to guard against malignant cells. The CD8^+^ T cell response to an antigen is composed of many T cell clones with unique T cell receptors, together forming a heterogenous repertoire of phenotypically and functionally distinct effector and memory cells^1, 2^. How individual T cell clones contribute to this heterogeneity during an immune response is key to understand immunity but remains largely unknown. Here, we longitudinally tracked CD8^+^ T cell clones expanding in response to yellow fever virus vaccination at the single cell level in humans. We show that only a fraction of the clones detected in the acute response persists as circulating memory T cells, indicative of clonal selection. Clones persisting in the memory phase displayed biased differentiation trajectories along a gradient of stem cell memory (SCM) towards terminally differentiated effector memory (EMRA) fates. Reactivation of single memory CD8^+^ T cells revealed that they were poised to recapitulate skewed differentiation trajectories in secondary responses, and this was generalizable across individuals for both yellow fever and influenza virus. Together, we show that the sum of distinct clonal differentiation repertoires results in the multifaceted T cell response to acute viral infections in humans.

## Main

To assess how individual clones together shape the heterogeneous repertoire of CD8^+^ T cells during acute and memory phases of an anti-viral T cell response in humans, we tracked thousands of antigen-specific CD8^+^ T cells longitudinally during primary immune responses to the Yellow Fever Virus vaccine YFV-17D. This is a very efficient, live, attenuated viral vaccine generating lifelong adaptive immunity following a single injection^3–7^. We monitored single YFV-17D-specific CD8^+^ T cells specific for the immunodominant HLA-A2 restricted epitope (NS4b LLW) as well as a subdominant HLA-B7-restricted epitope (NS5 RPI), during acute (day 15), early memory (days 90-136), and late memory (day 500+) phases of the response (Fig. 1a, Supplementary Figure 1a)^6^. We identified productive TCR*β* sequences for 3058 cells using a nested PCR strategy, targeting the TCR*β* chain locus in DNA from whole genome amplified single cells^8, 9^, and an additional 1720 cells using full length scRNA-seq and TCR*α* and/or *β* chain identification in four healthy human donors (Donors A-D) (Supplementary Table 1)^10, 11^. As previously reported^6^, the HLA-A2 response was an order of magnitude greater than the HLA-B7 response (Fig. 1b). Despite the difference in response magnitude, HLA-A2 and HLA-B7 responses displayed similar patterns of expansion and contraction (Supplementary Figure 1b, c). A significant drop in clonal diversity occurred during the transition from the acute (day 15) to memory phase of the response for all donors and epitopes (p<0.05) (Fig. 1c). Higher clonal diversity at day 15 was associated with increased frequency of responding cells (Fig. 1d), indicating that the number of distinct naïve precursor cells recruited to the response determines the response magnitude.

**Figure 1.**
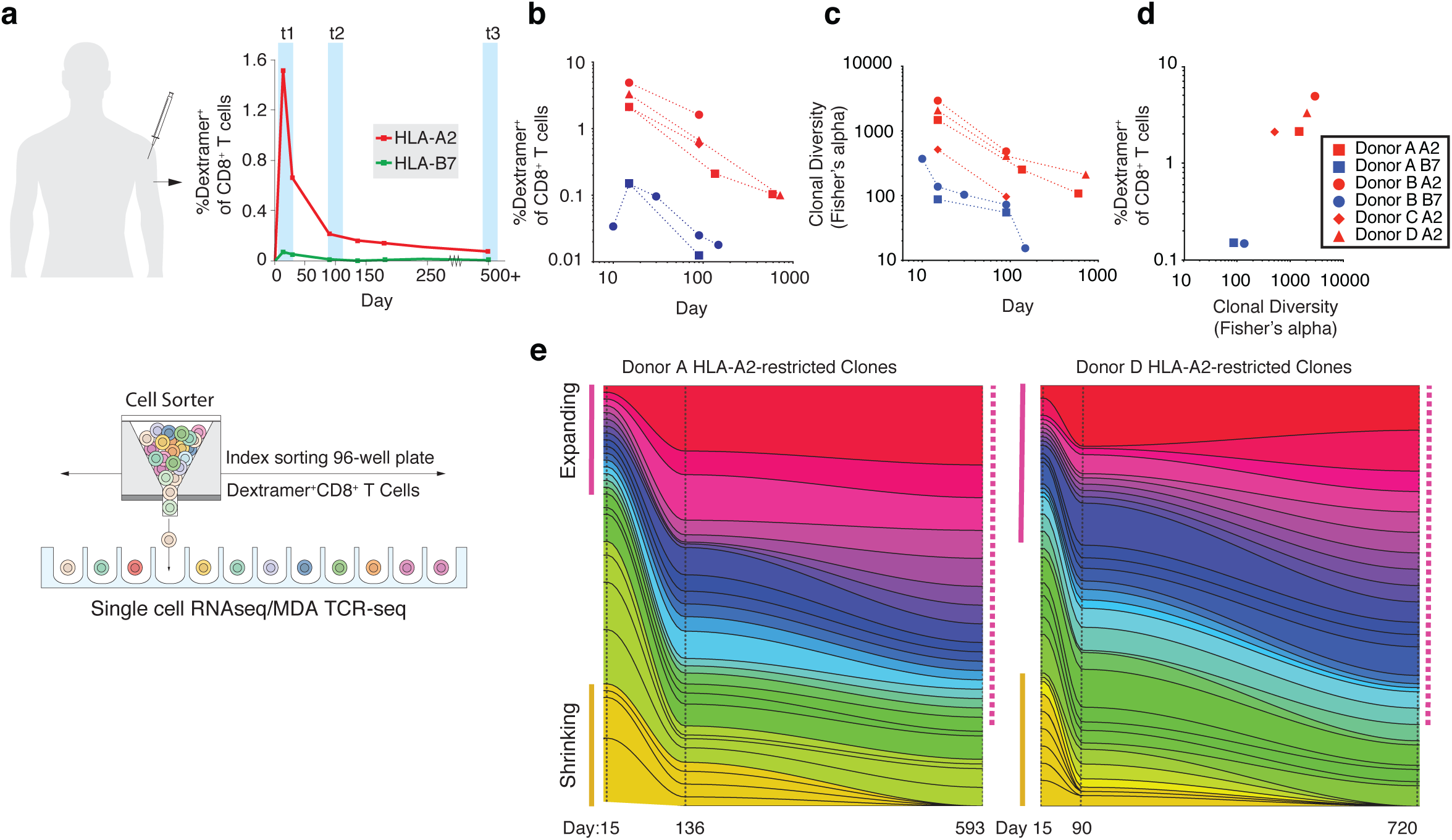
Clonal Contributions throughout the Primary Immune Response to Yellow Fever Vaccine. **a**, Schematic diagram for experimental setup. Frequency of HLA-A2/YFV- and HLA-B7/YFV-dextramer^+^ cells of total CD8^+^ T cells for donor A is shown as an example. **b**, Frequency of YFV-specific CD8^+^ T cells of total CD8^+^ T cells. (n=4 for HLA-A2, n=2 for HLA-B7) **c**, Clonal diversity measured by Fisher’s alpha throughout the acute to memory transition in donors A-D. **d**, Correlation between frequency of YFV-specific (dextramer^+^) CD8^+^ T cells versus clonal diversity (Fisher’s alpha) at day 15 (r^2^=0.89, p<0.01 by two-tailed Pearson). **e**, Evolution of clone sizes throughout the acute (day 15) to memory (days 90/136 and 593/720) transition identifies expanding clones which dominate the late memory response and shrinking clones which dominate the acute response. All HLA-A2-restricted clones from donor A and D with n>3 cells at a single timepoint are shown. A summary of statistical results can be found in table S2.

The decline in diversity over time indicates that clonal selection plays a role in shaping the circulating memory repertoire after YF-17D vaccination, consistent with focusing of the TCR repertoire observed in bulk analysis of cytomegalovirus (CMV) pp65-specific CD8^+^ T cells isolated from primary to chronic stages of CMV infection^12^. However, it is unclear whether expansion early in the response is an important determinant of selection into the memory repertoire. Adoptive transfer and lineage tracing studies in mice suggest that poorly expanded clones may preferentially persist as memory cells^13–15^, and extinction of some highly expanded clones has been observed in primary to chronic stages of CMV infection^12^, suggesting that not all highly expanded clones in the acute response transition into memory cells. Indeed, contraction of large clones and expansion of smaller clones was apparent during the acute to memory transition in two donors with datasets spanning the acute to late memory phases of the HLA-A2-restricted response (Fig. 1e). Analysis of the relationship between clone size and persistence to the late memory phase (day 593 or 720) revealed that a large clone size at day 15 was negatively associated with long-term survival (log odds ratio: −1.76 donor A, −1.83 donor D, p<0.05) while a small clone size at day 90/136 was positively correlated with persistence to day 500+ (log odds ratio: 0.51 donor A, 0.73 donor D, p<0.05) (Supplementary Table 2). Our results highlight the stability of the clonal repertoire throughout the memory phase of the immune response and demonstrate that large clonal expansion during the acute response is not a prerequisite for selection into the long-lived memory repertoire, and may instead preclude memory differentiation.

Diverse memory and effector lineages are known to persist for decades after YFV vaccination, and are a hallmark of memory T cell immunity^5, 16, 17^. Although classically these populations have been viewed as discrete cell types, high dimensional analysis of CD8^+^ populations increasingly suggest that these cells exist in a spectrum of cell types ranging from CCR7^+^ central memory (CM) and stem cell memory (SCM) to highly differentiated CCR7^−^ effector memory (EM) and terminally differentiated EM (EMRA)^2, 18^. We examined the phenotypic heterogeneity of the acute and memory response for CD8^+^ T cells specific for the HLA-A2/NS4b LLW and HLA-B7/NS5 RPI epitopes in three donors using high-parameter flow cytometry. We measured expression of proteins associated with CM/SCM (CCR7, CD127, TCF-1, LEF1, CD27)^17, 19–21^ and EM/EMRA (granzyme A, granzyme B, CD94, CD57)^22, 23^, in addition to CD45RA, KLRG1, CXCR4, and Ki67, and analyzed the heterogeneity using dimensionality reduction to clarify the distribution of YFV-specific CD8^+^ T cells across the spectrum of phenotypes over time^24^. The vast majority of acutely responding cells clustered together, between highly polarized effector memory and stem cell memory populations, and expressed effector proteins (granzymes A and B), yet were distinguished from memory populations on the basis of having lower expression of CD45RA and CXCR4 (Fig. 2a, Supplementary Figure 2a). By contrast, most cells at day 90 regained CD45RA expression and displayed increased phenotypic heterogeneity, covering the range of phenotypes between highly polarized SCM and EMRA fates (Fig. 2b, Supplementary Figure 2a). We noted clear interindividual differences in distribution of YFV-specific cells along the spectrum of memory identities (Supplementary Figure 2b, c), whereas no major differences were observed between HLA-A2 or HLA-B7 restricted viral epitopes (Supplementary Figure 2d). The analysis also revealed the existence of intermediate phenotypes, e.g. a population of CD127^+^CCR7^−^ CD8^+^ T cells with intermediate expression of TCF-1 and LEF-1 (Supplementary Figure 2e, f), further supporting the notion that CD8^+^ T cell differentiation encompasses a spectrum of identities rather than discrete cell types.

**Figure 2.**
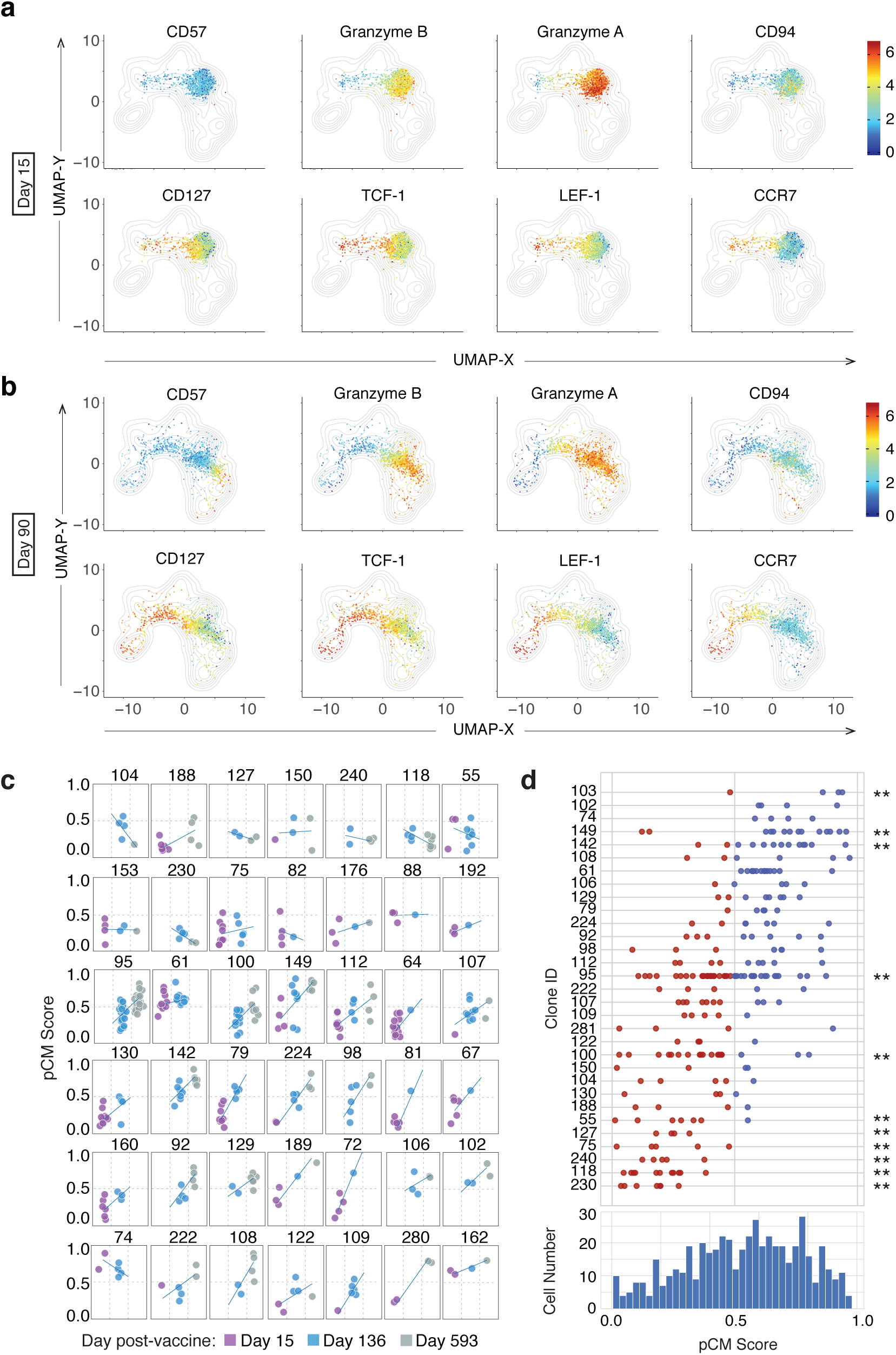
Emergence of Phenotypically Distinct Memory Subsets at the Population and Clonal Level following Primary YFV-17D vaccination. **a**, **b** Visualization of YFV-specific CD8^+^ T cell phenotypic distribution by UMAP dimensionality reduction based on protein expression during acute (**a**, day 15) or memory (**b**, day 90) phases of the primary response to YFV vaccination. Concatenated data from HLA-A2/YFV- and HLA-B/YFV-dextramer^+^ CD8^+^ T cells from three donors are shown. Bulk peripheral blood CD8^+^ T cells are included as contour plots in the background (gray). UMAPs were based on expression of CCR7, CD27, CD45RA, CD57, CD127, CD94, granzyme A, granzyme B, KLRG1, LEF-1, and TCF-1 in 8000 bulk CD8^+^ T cells and 2400 HLA-A2/YFV- and HLA-B7/YFV-dextramer^+^ CD8^+^ T cells concatenated from the three donors. Expression levels of individual protein markers are displayed as heatmaps (log2 scaled values according to color scale). Individual data is summarized in Supplementary Figure 2a. **c**, Individual T cell clones from donor A for all three time points plotted according to phenotype (pCM=0: strong EM/EMRA identity, pCM=1.0 strong CM/SCM identity). Only clones with n>3 cells found at least at 2 time points are shown (both HLA-A2 and B7-restricted clones are shown). **d**, Distribution of single cell phenotypes for clones during the memory (day 136/593) phase of the response according to pCM score. Clones with n>3 cells found at either timepoint for both HLA-A2 and B7-restricted responses are shown. Histogram represents the distribution of all cells in clones with n>3 cells along the gradient of pCM scores. ** Clones which significantly diverge from expected distributions based on total population (p<0.05 Wilcoxon Rank-Sum test).

Although our analysis of protein expression revealed that YFV-specific memory CD8^+^ T cells encompassed a range of phenotypes, it did not allow us to track the behavior of individual clones over time. To this end we combined cell surface protein expression analysis (Supplementary Figure 3a, b, Supplementary Table 3) with high coverage single-cell RNAseq^11^. This approach both allowed us to identify distinct clones and to further refine the classification of cellular marker profiles, coupling clonality to phenotype. Two healthy subjects were monitored throughout the acute to memory transition, and one donor was selected for in-depth clonal analysis. To quantify each cell’s identity along the continuum of phenotypes we used a recently described partial least squares regression approach, with CCR7 protein expression as a starting point, to train a classifier for all sequenced YFV-specific CD8^+^ T cells (Supplementary Figure 4a)^25^. This allowed us to rank and assign a score to each CD8^+^ T cell based on transcriptome differences according to the likelihood that they were strong effector (pCM=0) or strong SCM cells (pCM=1.0) (Supplementary Figure 4b-d). We validated the pCM score on cells sorted from bulk CD8^+^ T cell populations on the basis of classically defined phenotypes (Naïve, SCM, CM, EM, EMRA) (Supplementary Figure 5). Notably, many of the top genes identified by our classification strategy (*TCF7*, *CCR7*, *LTB*, *SELL*, *GZMK*, *GZMB*, *GZMH*, *ZEB2*, *CCL4*, *GNLY*) play established roles in specifying SCM and effector memory cell identities in mice and humans^2, 20, 26, 27^. In both donors we found that the pCM score accurately ordered cells according to expression of these genes (Supplementary Figure 4b-d). For Donor A, for which three timepoints were collected, we analyzed how individual clones varied with respect to pCM score throughout the response (Fig. 2c). In the acute phase of the response, most clones had low pCM scores, indicating an effector bias. In the early and late memory phase, we observed a gradual increase in pCM scores over time for cells in many of the clones, consistent with the idea that some clones transition via an effector stage to an SCM fate^3^. Focusing on the memory phase we found that not all clones, however, transitioned towards a SCM fate, as some gave rise exclusively to effector cells (Fig. 2c, d). Our findings thus reveal that distinct clonal fates in the memory phase of the response give rise to the overall phenotypic heterogeneity observed in long-lived CD8^+^ T cell memory repertoires.

**Figure 3.**
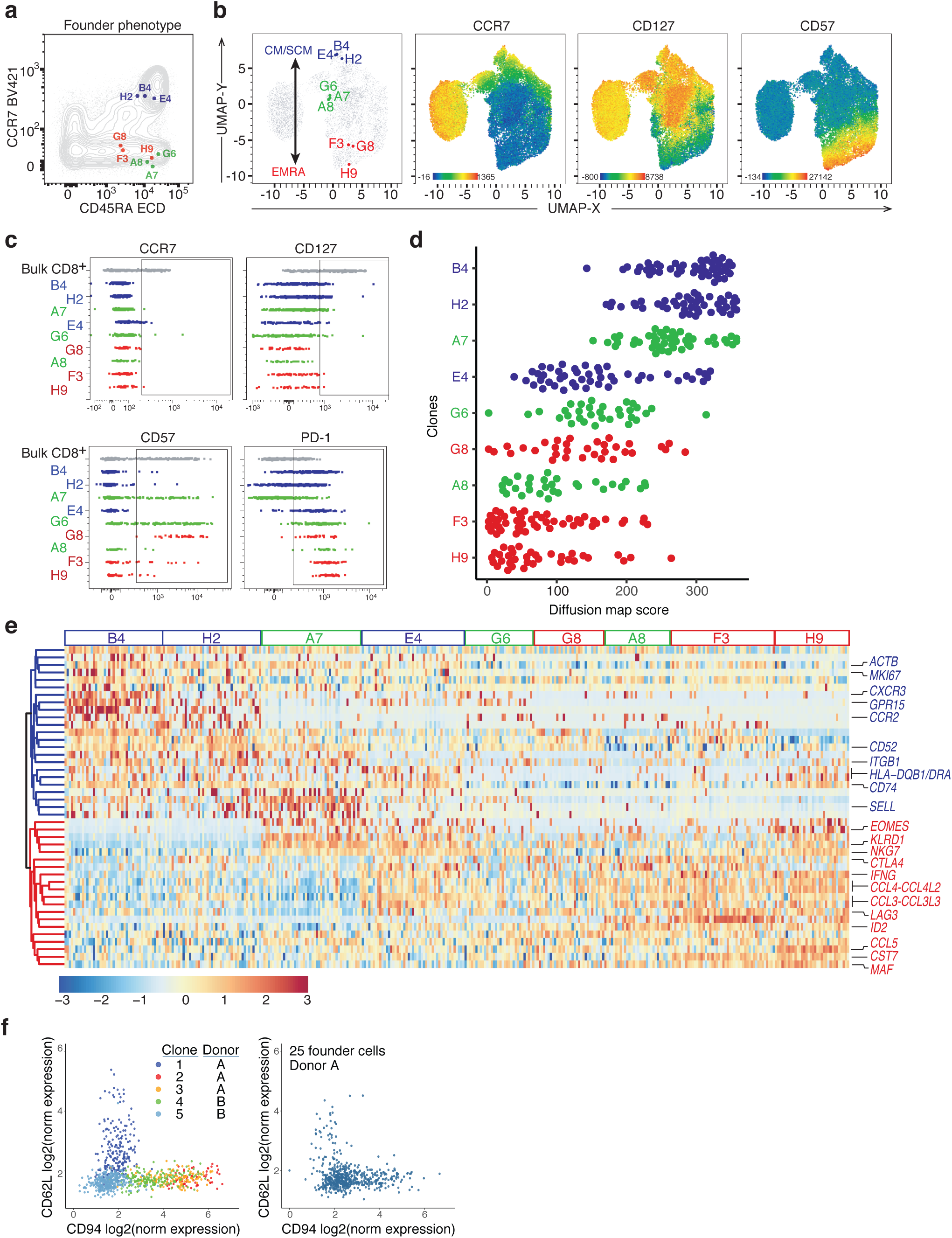
Secondary Reactivation of Single CD8^+^ T Cell Clones Yields Phenotypically Distinct Effector Progeny. **a**, Phenotype of single, index-sorted CD8^+^ T cell clones from Donor A at Day 136 (founder phenotype) overlaid on bulk CD8^+^ T cells from matched blood, using classical EM/EMRA and CM/SCM classification (CD45RA vs CCR7, contour plot). **b**, UMAP dimensionality reduction plots depicting bulk CD8^+^ T cells (gray) and individual sorted SCM (CCR7^+^CD127^+^; blue), CD127^+^ EMRA (CCR7^−^CD127^+^CD57^−^; green), and CD127^−^ EMRA (CCR7^−^CD127^−^CD57^+^; red) YFV-specific founder CD8^+^ T cells from Donor A (day 136) (far left). Color gradient in UMAPs indicate protein expression levels for CCR7, CD127, and CD57 on total CD8^+^ T cells. UMAPs were based on expression of CCR7, CD3, CD8, CD27, CD45RA, CD57, CD95, CD127, and PD-1. **c**, Cell surface protein expression of CCR7, CD127 (SCM/CM markers), CD57 and PD-1 (effector/activation markers) on expanded T cell clones (B4-H9) from each sorted single founder T cell in (**a**) after 20 days of stimulation with YFV-peptide (HLA-A2/LLWNGPMAV) and IL-2 in the presence of irradiated autologous CD3-depleted PBMC. Unstimulated bulk CD8^+^ T cells from peripheral blood are depicted at the top of each graph in gray as a reference. **d**, Diffusion map score (using DM1) for expanded single T cells from each clone based on variable genes identified for single cell transcriptomes from each clone (n=32-45 cells/clone). **e**, Heatmap depicting a selection of differentially expressed genes based on analysis of expanded single cells from CCR7^+^ (B4, H2, E4) versus CCR7^−^ (A7, G6, G8, A8, F3, H9) founders. Genes clustered according to Euclidean distance. Genes are colored according to enrichment in expanded T cell clones originating from SCM (blue) or EMRA (red) founders. **f**, CD94 and CD62L protein expression on individual clones expanded *in vitro* for 20 days from founders isolated from donor A (day 136 after vaccination) and donor B (day 90 after vaccination). Right panel shows phenotype of progeny from 25 sorted memory T cells stimulated in a single well from donor A (day 136 after vaccination).

**Figure 4.**
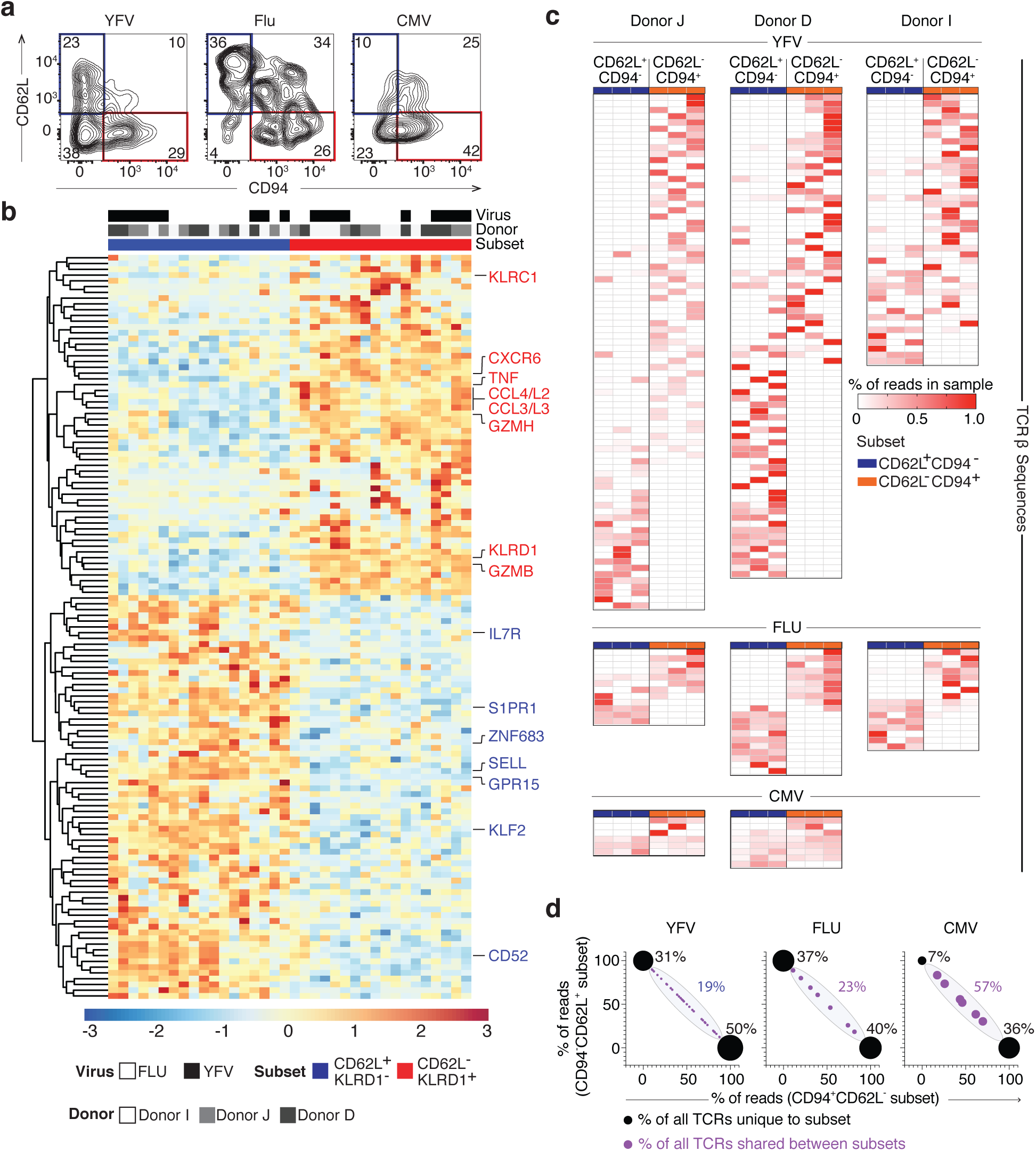
Clonally Distinct Effector Populations emerge after Secondary Activation of bulk YFV- and Influenza-Specific Memory CD8^+^ T cells. **a**, Expression of CD62L and CD94 on expanded virus-specific (HLA-A2/virus-dextramer^+^CTV^−^) CD8^+^ T cells 2-weeks after stimulation with YFV, Influenza or CMV from a single donor (Donor I; Donors J and D in Supplementary Figure 9b). **b**, Heatmap depicting differentially expressed genes identified by RNA-seq (padj<0.05, log2-fold change>1) between CD62L^+^CD94^−^ and CD62L^−^CD94^+^ expanded YFV- and influenza-specific CD8^+^ T cells from three donors (3 replicates with 50 cells each per donor/virus combination). Scale represents z-score and virus and donor are depicted in top bars. Euclidean clustering segregates effector T cells on the basis of phenotype, but not on the basis of donor origin or viral specificity. Heatmap for CMV-specific CD8^+^ T cells is shown in Supplementary Figure 9c. **c**, Heatmaps depicting distribution of unique TCRβ chains in different subpopulations for each donor/virus combination identified from bulk RNAseq replicates. Each row represents a unique TCRβ chain. Triplicates with 50 cells for each donor/virus/ subpopulation were used to estimate clonal abundance in each group. TCR sequences which were observed across different viral responses within a single donor were excluded from the analysis to remove false positives. **d**, Distribution of TCRβ chains between CD62L^+^CD94^−^ and CD62L^−^CD94^+^ subsets in YFV, influenza, and CMV-responding CD8^+^ T cells. Y- and X-axes represent percent of reads found in CD62L^+^CD94^−^ and CD62L^−^CD94^+^ subsets respectively. Size of circles and percentages represent percent of all clones found in either the CD62L^+^CD94^−^ or the CD62L^−^CD94^+^ subsets (black filled circles) or that are shared between subsets (blue filled circles).

Secondary infections result in a rapid mobilization of memory T cell subsets to produce a diverse population of effector cells. The current model postulates that CM/SCM act as a pluripotent, self-renewing population which gives rise to all effector cell classes ^21, 28, 29^, whereas EM/EMRA only are capable of short bursts of highly cytotoxic effector cells^17, 30, 31^. However, it is not possible to infer properties at the single cell level from bulk analysis. To address how different memory T cell types respond at the single cell level, we performed antigen-specific reactivation of sorted, individual YFV-specific memory CD8^+^ T cells isolated from the memory population (day 136) and assessed the phenotype and heterogeneity of their clonal progeny by flow cytometry and scRNAseq. We characterized the response of 9 unique clones identified as SCM (clones B4, E4, H2), CD127^+^ EMRA (clones A7, A8, G6) or CD127^−^ EMRA (clones F3, G8, H9) which represent the range of memory identities observed in our study (Fig. 3a, b). After a 20-day expansion period we detected thousands of progeny from each clone, all of which were CCR7^−^CD127^−^ and which expressed variable levels of PD-1 and CD57, indicating their status as highly activated effectors (Fig. 3c). Notably we observed higher PD-1 expression in progeny derived from EMRA compared to SCM (Fig. 3c), suggesting a stronger predisposition of EMRA-derived progeny to develop senescent or ‘exhausted’ phenotypes in recall responses^32^. In contrast to EMRA-derived clones, SCM-derived and CD127^+^ EMRA-derived T cell clones displayed signs of proliferation even 20 days post-activation (Supplementary Figure 6a), in line with the view that SCM cells undergo greater expansion in recall responses^21^.

We performed scRNAseq on 36-45 cells in the progeny from each clone to define the phenotypic heterogeneity at the single cell level within and between clones. After accounting for cell cycle differences, we examined intra- and inter-clonal variability of effector progeny according to gene expression differences by diffusion map dimensionality reduction (Fig. 3d)^33^. This revealed distinct clonal gene expression signatures of effector progeny derived from different founder cells. We observed that clonal effector progeny generally separated on the basis of founder cell phenotype, albeit with cells once again ordering along a continuum of identities as opposed to discrete cell types. There were however exceptions; clone E4, which had a clear SCM founder phenotype, produced progeny more similar to progeny from CD127^+^ EMRA founders, while clone A7, a clone derived from a CD127^+^ EMRA progenitor, produced progeny similar to those from SCM founders (Fig. 3d). The observation that different memory T cell clones could yield distinct effector progeny, despite exhibiting stereotypical SCM phenotypes, also provides a potential explanation for how heterogeneous secondary responses can be generated after reactivation of bulk SCM populations^19, 21^. Analysis of differentially expressed genes between progeny from SCM and EMRA founders identified an enrichment of classical effector genes (*IFNG*, *CCL3*, *CCL4*, *CCL5*) in EMRA-derived progeny, while there was an enrichment of genes involved in lymphocyte homing (*CXCR3*, *GPR15*, *ITGB1*, *CCR2*, *SELL*) in SCM-derived progeny (Fig. 3e, Supplementary Figure 6b, c, Supplementary Table 4).

We identified *SELL* (CD62L) and *KLRD1* (CD94) as cell surface markers which generally segregated SCM- and EMRA-derived progeny. In additional experiments we observed that progeny of single activated clones often expressed either CD62L or CD94 (Fig. 3f). Comparison of individual clonal recall responses by mixed (bulk, 25 memory cells) YFV-specific cells from one donor isolated 1401 days post-vaccination, demonstrated that the mixed effector profiles observed after secondary stimulation were still observed over 3 years into the memory phase of the response (Supplementary Figure 7a-c).

Based on our findings, we propose a model in which the long-lived memory repertoire consists of clones with divergent phenotypes, that give rise to a range of distinct effector cell types after reactivation (Supplementary Figure 8). To test whether divergent clonal phenotypes was a generalizable phenomenon shared also in recall response to other viruses, we designed an experiment to measure responses by memory CD8^+^ T cell populations against an additional acute infection (Influenza) and a chronic infection, CMV, in parallel to responses to YFV. We identified three healthy HLA-A2^+^ individuals with detectable levels of both HLA-A2/YFV- and HLA-A2/Influenza (M158–66; GILGFVFTL)-specific CD8^+^ T cells, two of which also had detectable levels of CMV (pp65; NLVPMVATV)-specific CD8^+^ T cells. We labeled total peripheral blood mononuclear cells (PBMCs) from each donor with CellTrace violet (CTV) and stimulated one million PBMCs with each viral peptide in parallel. After two weeks we observed the presence of expanded, antigen-specific CD8^+^ T cells for each viral antigen (Supplementary Figure 9a). We phenotyped the expanded T cells for each donor/virus combination using CD62L and CD94 expression and analyzed triplicates of 50 sorted CTV^−^ Dextramer^+^ CD8^+^ T cells from CD62L^+^CD94^−^ and CD62L^−^CD94^+^ subsets using RNAseq (Fig. 4a, Supplementary Figure 9b). Differential gene expression analysis of CD62L^+^CD94^−^ versus CD62L^−^CD94^−^ CD8^+^ T cells specific for YFV and Influenza revealed a clear segregation of the two populations across donors and anti-viral responses (Fig. 4b, Supplementary Table 5)). Gene expression differences were similar to our clonal scRNAseq experiments, with CD62L^+^CD94^−^ cells expressing elevated levels of genes related to homing (*GPR15*, *S1PR1*, *KLF2*) and CD62L^−^ CD94^+^ cells expressing elevated levels of effector-associated genes (*CCL3*, *CCL4*, *CCL5*, *TNF, NKG7*) (Fig. 4b). In contrast to the YFV and influenza-specific CD8^+^ T cells, we detected substantially less differences between CD62L^+^CD94^−^ and CD62L^−^CD94^+^ CMV-specific CD8^+^ T cells, possibly reflecting a larger overlap in effector identities in the setting of chronic viral exposure (Supplementary Figure 9c).

To determine whether the CD62L^+^CD94^−^ and CD62L^−^CD94^+^ populations from each virus-specific CD8^+^ T cell population also had distinct clonal compositions we analyzed the distribution of TCR*β* sequences between samples (Fig. 4c, Supplementary Table 6). YFV-specific responses were characterized by a higher number of unique clones relative to influenza and CMV specific responses in all donors (Fig. 4c). The CD62L^+^CD94^−^ and CD62L^−^CD94^+^ populations, in both YFV- and influenza-specific effector cells, largely contained unique and non-overlapping clones (Fig. 4c, d), demonstrating that averaging of distinct clonal differentiation trajectories contributes to the phenotypic heterogeneity in recall responses in both YFV and influenza infection. In contrast to YFV and influenza, CMV responses were characterized by substantial clonal overlap between the two populations (Fig. 4c, d), suggesting that chronic antigen exposure can lead to subclonal diversification and thus increased clonal heterogeneity, or possibly promote the differentiation of polyfunctional clones.

Different models have been proposed to explain the existence of phenotypically distinct memory T cell subsets after infections^34^. Here we demonstrate that clonal selection after the acute phase shapes the repertoire of circulating memory CD8^+^ T cells, leaving clones undergoing biased differentiation trajectories towards distinct memory phenotypes along the continuum of SCM to EMRA identities. Clonally biased differentiation fates were also observed after reactivation at the single level, suggesting that different memory T cell clones play distinct roles in secondary immune responses. Recent evidence suggests that circulating effector cells may act as precursors for tissue-resident memory CD8^+^ T cells^35^. It is therefore tempting to speculate that the clonal division of labor observed here could translate to a spatial segregation of clonal responses, where activation of multiple spatially distinct T cell clones would be required to mount the full repertoire of CD8^+^ T cells in secondary responses. Focusing future efforts on tracking the fate and function of T cells at the clonal level should provide valuable insights into the mechanisms determining the generation of memory T cell diversity, and perhaps more importantly how clonal identity impacts secondary immune responses, which may guide the development of new vaccines.

## Acknowledgements

We thank the Sequencing Core Facility at SciLife Labs for help with sequencing and data management. M. Toro and S. Giatrellis provided assistance with FACS.

## Author Contributions

J.E.M., J.M., and J.F. designed the study and wrote the paper. J.E.M. and J.M. performed all experiments. J.E.M., J.M., L.M.M., J.H. M.Z., A.J.M.L., C.J.E., P.S., E.B., C.T.L., B.A. P.R., and F.P. were involved in data analysis and provided support for data analysis. J.M., K.B. and J.S collected samples and data relevant to YFV-17D longitudinal study. B.R., S.P. and R.S. provided assistance with single cell RNA-seq studies.

## Data availability statement

All sequence data that support the findings of this study will be deposited at the European Genome-Phenome Archive (EGA) before publication of the manuscript.

## Code availability statement

All code used for analysis has previously been published and is available.

---

**Supplementary Figure 1.**
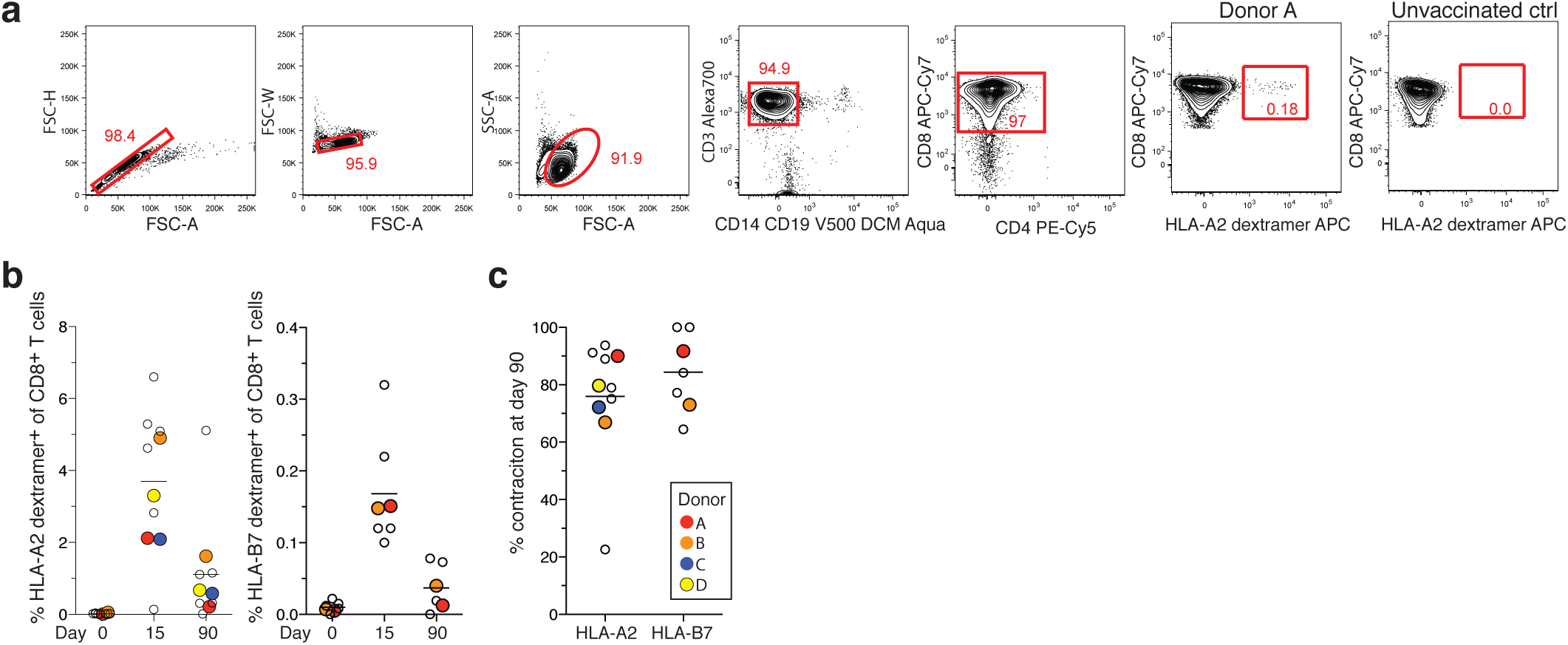
Sorting gates and frequencies of YFV-specific CD8^+^ T cells found at different times post-vaccination. **a**, Single live, CD14^−^ CD19^−^ CD8^+^Dextramer^+^ T cells were sorted for single cell RNAseq (Donor A shown). Background staining for HLA-A2/YFV-dextramer in an unvaccinated healthy donor shown as control. **b**, Frequency of HLA-A2/YFV (left) and HLA-B7/YFV (right) dextramer^+^ CD8^+^ T cells at days 0, 15, and 90 after vaccination. **c**, Percent contraction of CD8^+^ T cell responses to HLA-A2/YFV and HLA-B7/YFV from day 15 to day 90 for different donors (HLA-A2/YFV n=10, HLA-B7/YFV n=7). Colored circles represent donors included in single cell TCR analysis in main Fig. 1 and 2. Frequency data for all donors except donor A and B were sourced from Blom et al (6).

**Supplementary Figure 2.**
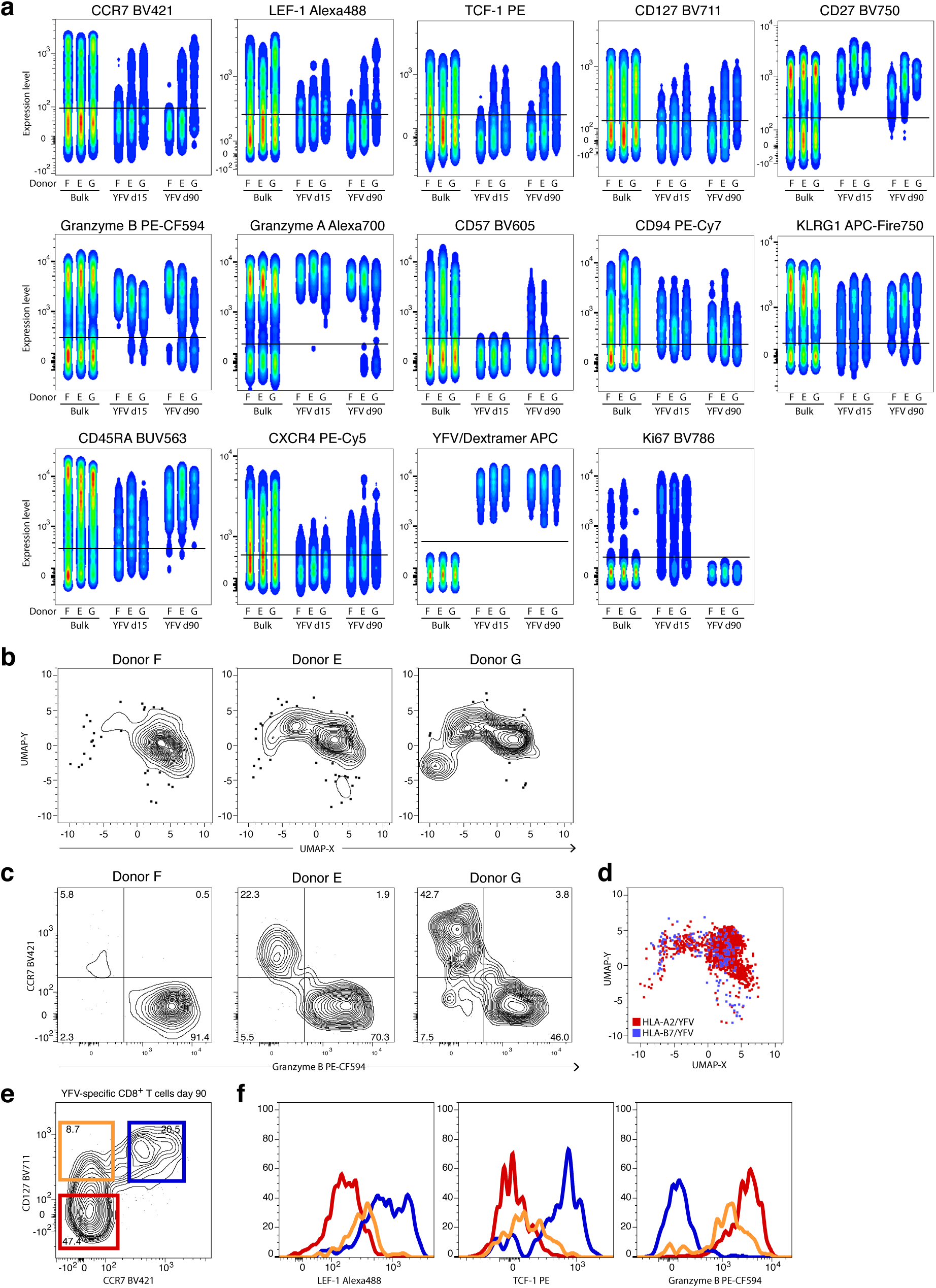
Donor and time-dependent changes in phenotype of YFV-specific CD8^+^ T cells. **a**, Protein expression by bulk CD8^+^T cells and YFV-specific CD8^+^ T cells at d15 and d90 analyzed by multicolor flow cytometry in three donors (donors E, F, and G) **b**, UMAP projection of protein expression data in three individual donors. **c**, flow cytotmetry analysis of CCR7 and granzyme B protein expression by YFV-specific CD8+ T cells at d90 after vaccination for three individual donors reveal donor-biased CD8^+^ T cell differentation fates. **d**, Overlay of UMAP projections of HLA-A2 (red) and HLA-B7 (blue) YFV-specific CD8^+^ T cells. **e**, Flow cytometry analysis of CD127 and CCR7 expression at day 90 after vaccination reveals an intermediate CCR7^−^CD127^+^ population (green). Data depict concatenated data from three donors. **f**, LEF-1 (left), TCF-1 (middle), and granzyme B (right) expression by CCR7^−^CD127^−^ (red), CCR7^+^CD127^+^ (blue), and CCR7^−^CD127^+^ (green) YFV-specific CD8^+^ T cells at day 90 after vaccination illustrates the continuous nature of CD8^+^ T cell differentiation states. Data depict concatenated data from three donors.

**Supplementary Figure 3.**
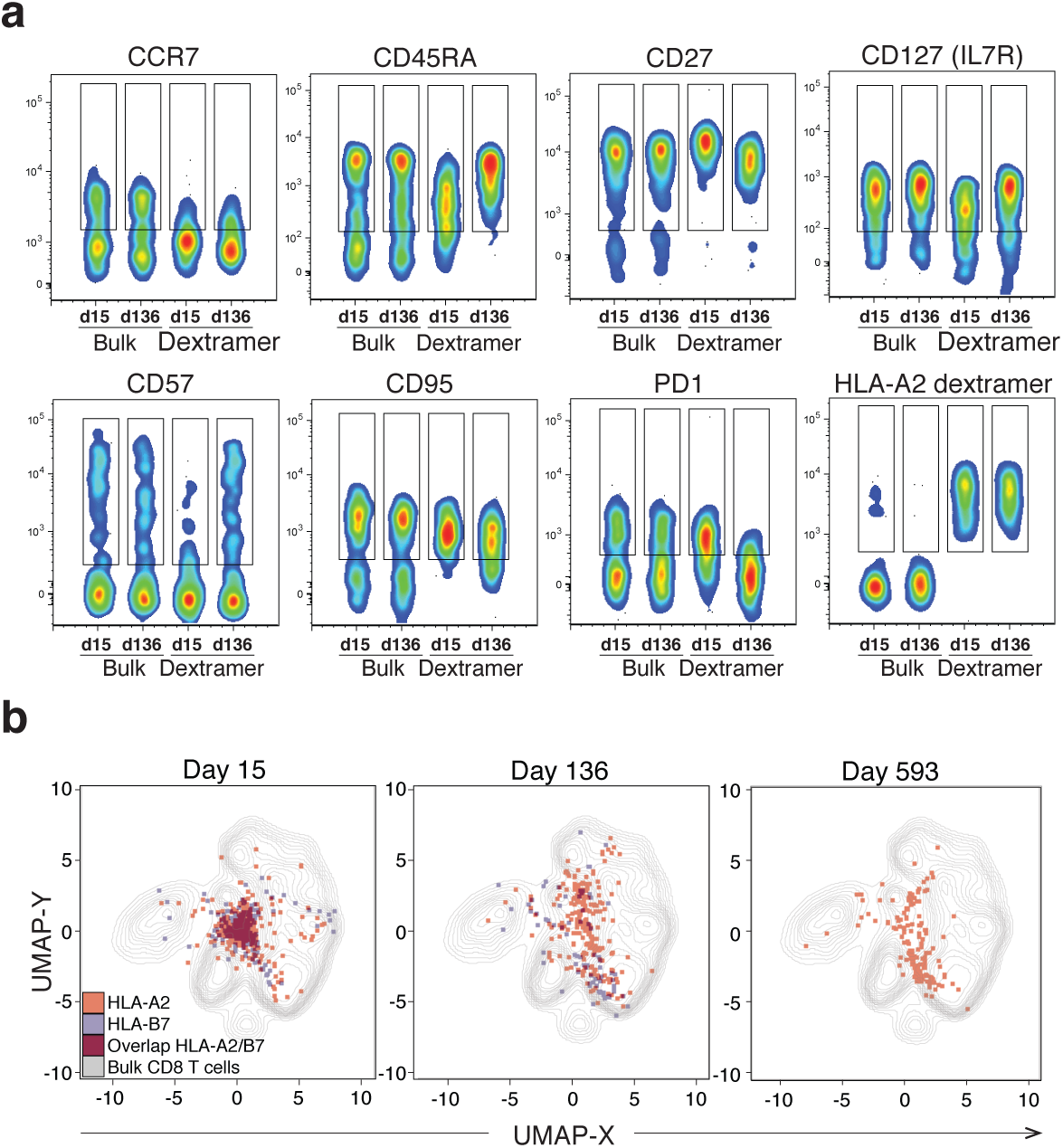
Cell surface phenotype of bulk and YFV-specific CD8^+^ T cells index-sorted for single cell RNAseq analysis. **a**, Protein expression by bulk CD8^+^ T cells and YFV-specific CD8^+^ T cells at day 15, 136, and 593 analyzed by multicolor flow cytometry. **b**, Projection of HLA-A2/YFV-specific (red) and HLA-B7/YFV-specific (blue) CD8^+^ T cells on UMAP based on expression of CCR7, CD45RA, CD27, CD127, CD57, CD95 and PD-1. Note large overlap in phenotype between HLA-A2/YFV- and HLA-B7/YFV-specific CD8^+^ T cells (depicted as dark grey). Bulk CD8^+^ T cells are shown as grey contour plot. Donor A is shown.

**Supplementary Figure 4.**
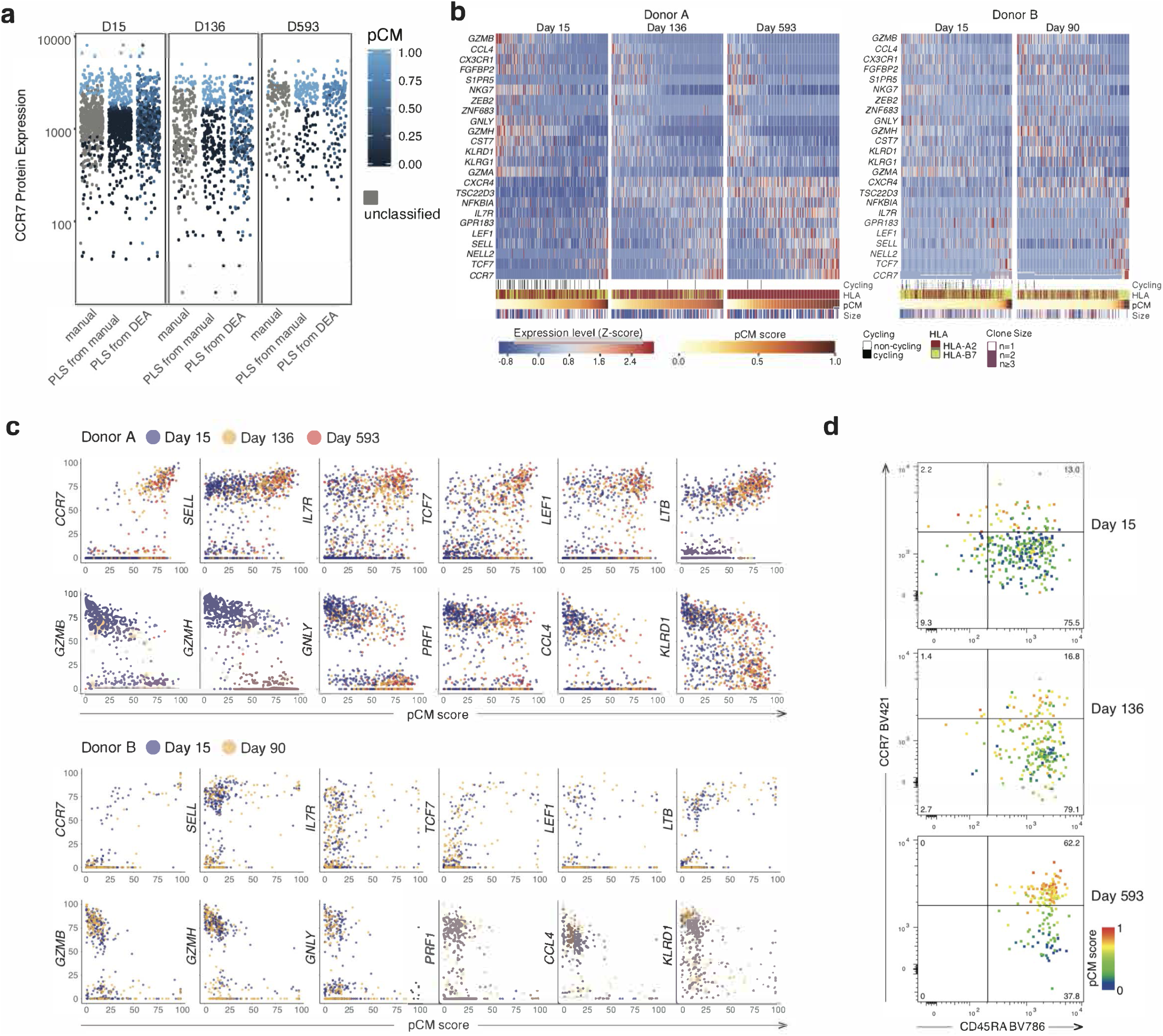
I Partial least square (PLS) analysis allows scoring of individual YFV-specific cos^+^ T cells along a continuum of cell states. **a**, Strategy for scoring YFV-specific CD8+ T cells by PLS. 101 cells with the highest and lowest CCR? protein expression respectively were chosen as starting point for PLS (manual, left)), and were ordered according the result from the PLS (PLS on manual, middle), and finally ranked based on results of PLS based on expression of differentially expressed genes between cells defined as effectors (pCM<0.5) and central memory (pCM>0.5) by PLS on manual. Detailed information about the analysis can be found in material and methods section. Donor A is shown. **b**, Heatmaps depicting gene expression along the pCM continuum at day 15, 136 and 593 for donor A, and at day 15 and 90 for donor B. **c**, Expression of canonical central and effector memory genes versus pCM score in donor A and donor B. **d**, pCM score in YFV-specific cells at days 15, 136, and 593 after vaccination in donor A depicted as color-scale relative to protein expression of CCR? and CD45RA.

**Supplementary Figure 5.**
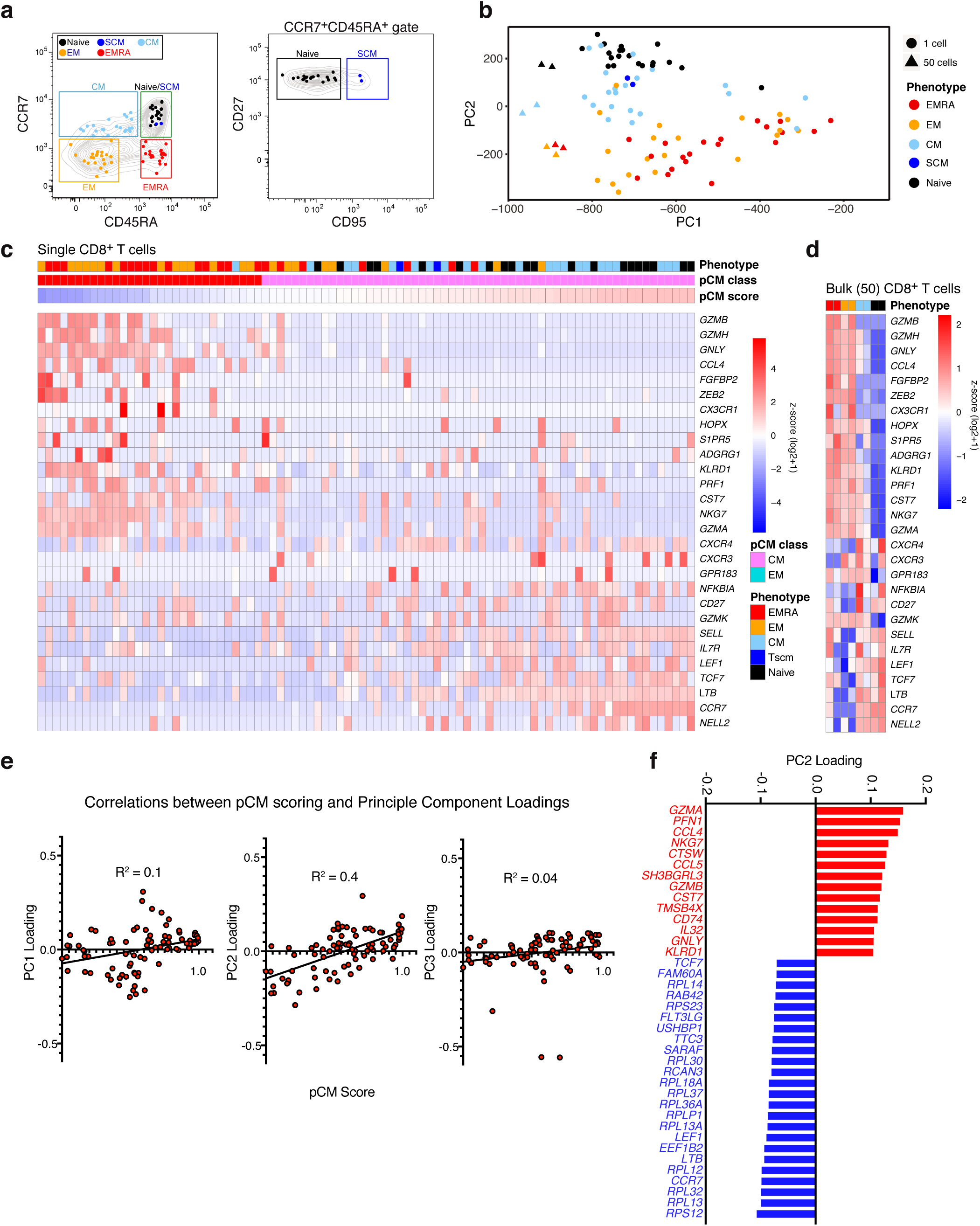
Application of pCM scoring strategy to single CD8^+^ T cells isolated from blood on the basis of classical phenotypic markers highlights continuous nature of CD8^+^ T cell subsets and accuracy of PLS method. **a**, Gating strategy to identify naïve (black), central memory (CM; light blue), effector memory (EM; orange), and terminally differentiated memory (EMRA; red) cells in bulk CD8^+^ T cells in peripheral blood. Two stem cell memory (SCM; dark blue) cells were ideintified among CD45RA^+^CCR7^+^ cells based on CD95 expression. Single cell RNA sequencing was performed on each cell, in addition to bulk (50 cells) from each of the naïve, CM, EM, and EMRA populations **b**, Principle component analysis of single and bulk CD8+ T cells from each population separates distinct subsets on PC2. Note that EM and EMRA cluster together, separately from naïve and CM CD8^+^ T cells. **c**, Expression of selected genes in single cells ordered according to pCM score. **d**, Expression of selected genes in bulk (50 cells) ordered according to cell surface phenotype. **e**, Correlation between pCM score and PC loadings shows that PC2 loadings correlate significantly with pCM score. **f**, Top genes contributing to PC2 loadings.

**Supplementary Figure 6.**
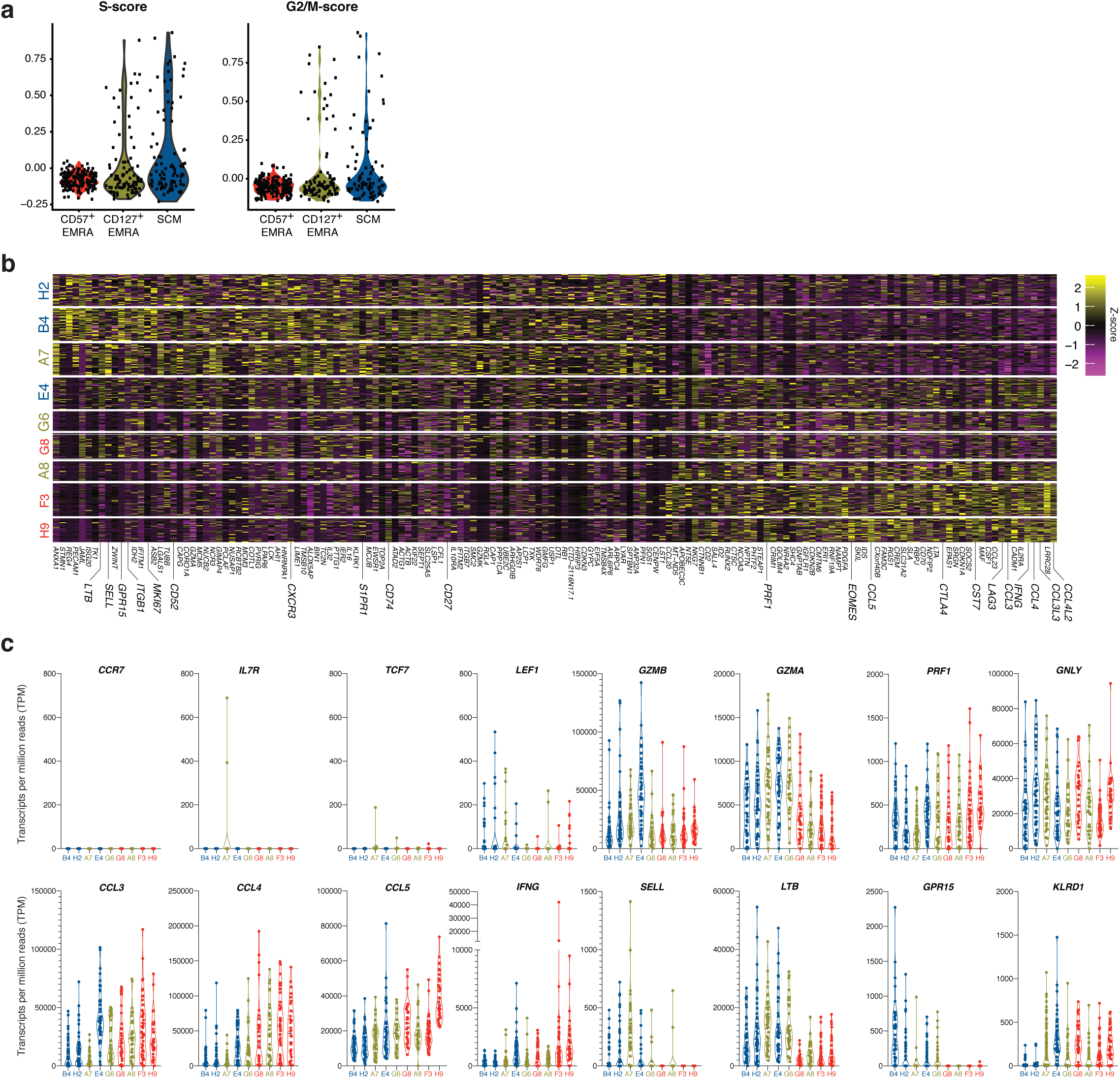
Effector progeny of single *in vitro* expanded YFV-specific memory CD8^+^ T cell clones have distinct phenotypes. **a**, Cell cycle analysis of single cell RNAseq data reveals that progeny from SCM and CD127**^+^** EMRA founders contain more dividing cells compared to CD57**^+^** EMRA founders even after 20 days of expansion in vitro. Cell cycle analysis was carried out using Seurat. **b**, Heatmap depicting differentially expressed genes between progeny of SCM and EMRA founder cells. **c**, Expression of selected genes associated with memory and effector CD8**^+^** T cells reveals that progeny from all clones display characteristics of effector cells, including lack of *CCR7*, *IL7R*, and *TCF-7*, and expression of *GZMA*, *GZMB*, *PRF1* and *GNLY*, yet are distinguishable by expression of chemokines and homing receptors.

**Supplementary Figure 7.**
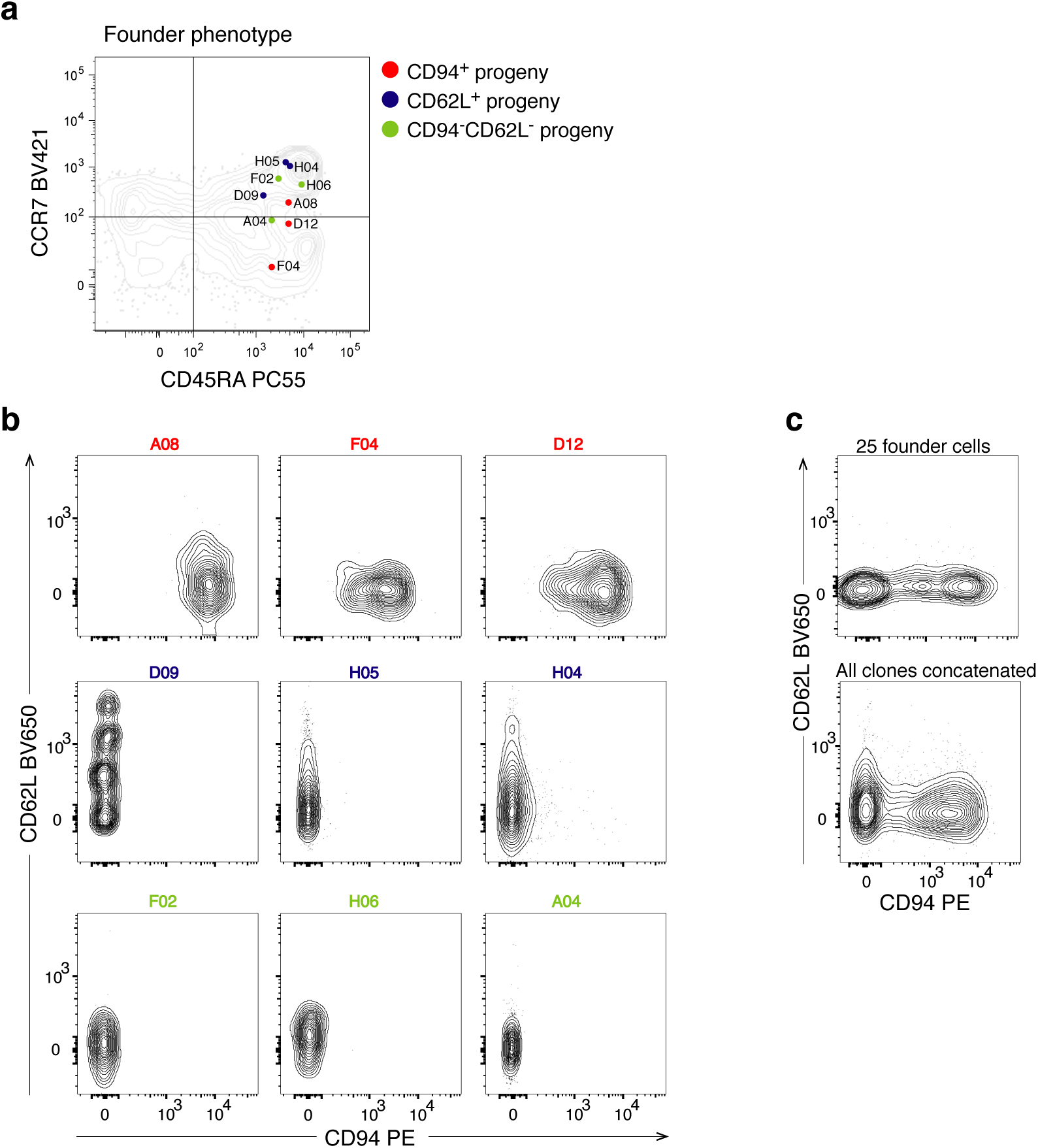
Secondary clonal effector responses by long-lived memory-founder CD8^+^ T cells give rise to heterogeneity through averaging of clonal identities. **a**, Founder cell identity related to expression of CCR7 and CD45RA protein expression. Founder cells are colored according to the phenotype of their subsequent progeny. Founder cells were sorted from donor B at d1400 after vaccination and expanded for 20 days with YFV NS4b LLW peptide, 50U IL-2/ml and irradiated autologous feeders. **b**, Expression of CD94 and CD62L on clonal progeny from (a). **c**, CD94 and CD62L expression in cultures where 25 founders were co-cultured (top) and concatenated data from the progeny of the 9 individual clones (bottom).

**Supplementary Figure 8.**
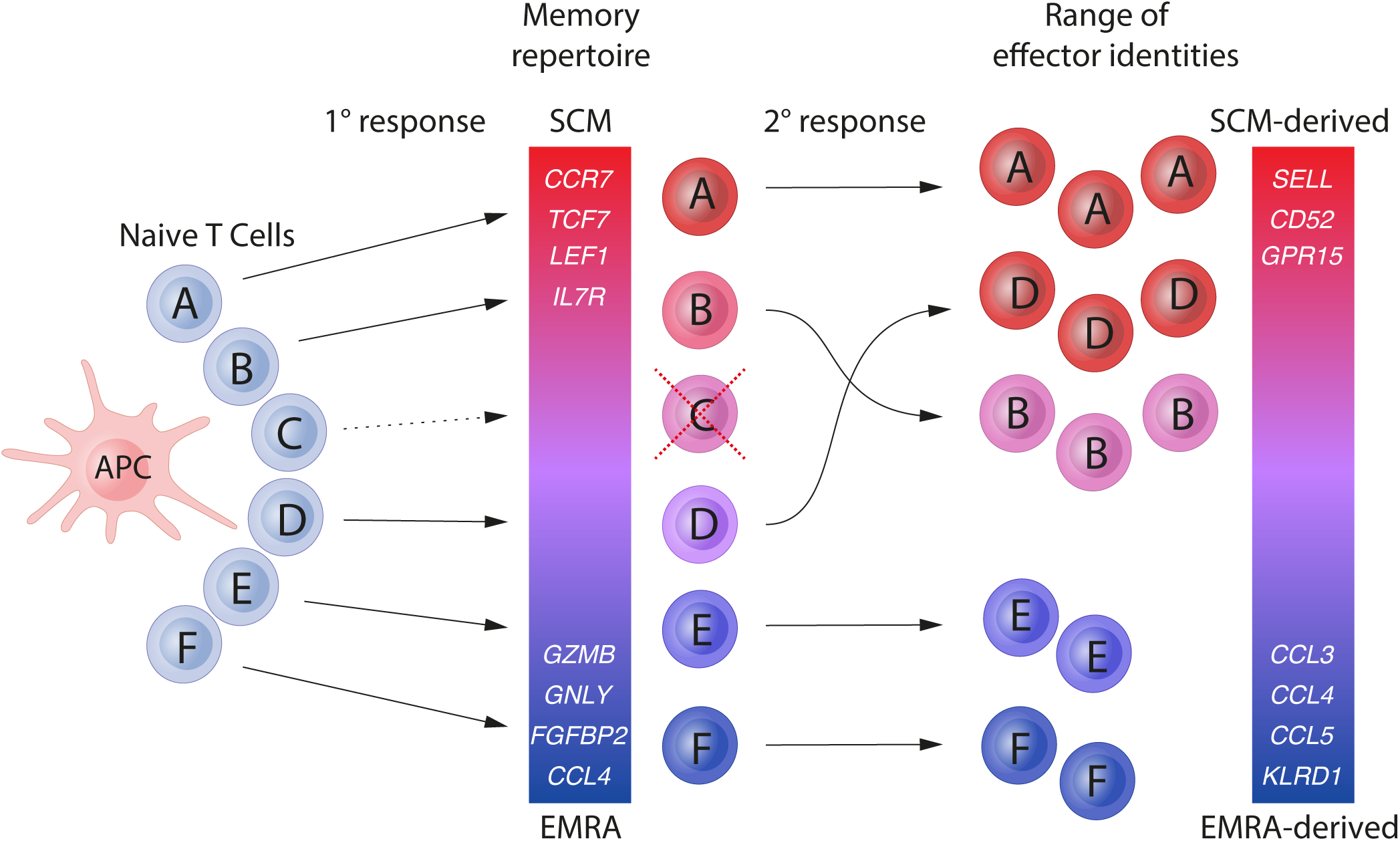
Distinct clonal identities in primary memory CD8^+^ T cells and in secondary recall responses. After vaccination a number of unique naïve CD8^+^ T cells recognize HLA class I/YFV-peptide complexes on the surface of antigen presenting cells, leading to massive clonal expansions. Some clones persist throughout the memory phase of the response, and exhibit biased fate trajectories (clones A, B, D, E, F), whereas others are no longer detectable at this stage despite large expansion in the acute phase of the response (clone C). Together, the distinct clonal identities combine to form the heterogenous repertoire of memory CD8^+^ T cells. Upon rechallenge with the antigen (2° response), memory CD8^+^ T cells are activated and expand, producing a range of phenotypically distinct effector CD8^+^ T cell types which vary between clones but tend to be similar within any given clone. In general SCM-derived progeny proliferate more and produce progeny with a bias towards expressing genes related to homing/adhesion (e.g. SELL, CD52, GPR15), whereas EMRA-derived progeny are biased towards expression of chemokines (CCL3, CCL4, CCl5).

**Supplementary Figure 9.**
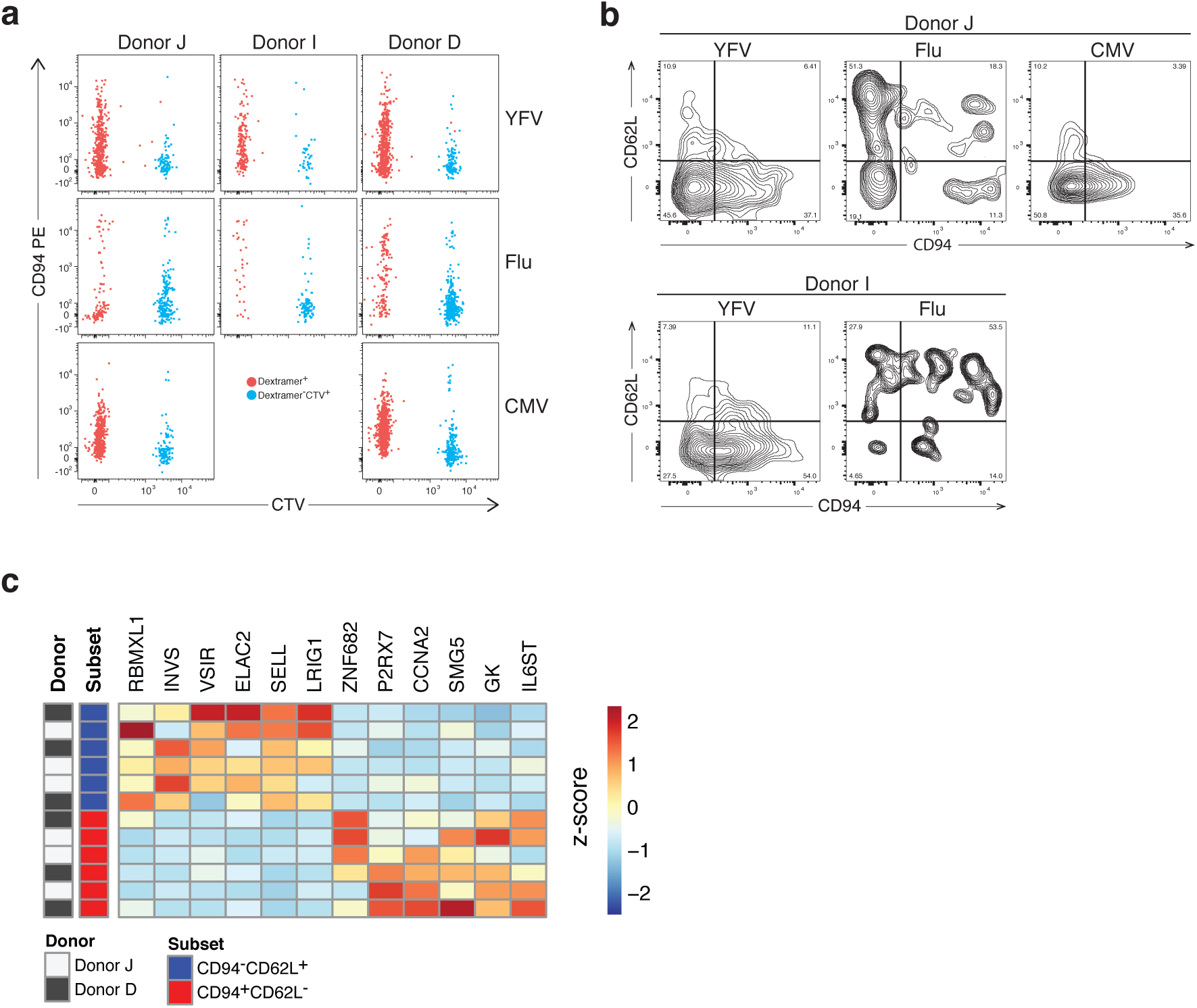
Expansion of YFV-, Flu-, and CMV-specific CD8^+^ T cells yields clonally distinct populations of CD94^+^CD62L^−^ and CD94^−^CD62L^+^ progeny. **a**, CellTrace Violet (CTV) dilution of YFV-, Flu-, and CVM-specific CD8^+^ T cells two weeks after stimulation with YFV LLW, Flu GIL, or CMV pp65 peptides and 50U IL-2/ml. Dextramer^+^ CD8^+^ T cells are shown in red, and bulk CTV^+^ CD8^+^ T cells are shown in blue. b, Expression of CD62L and CD94 on YFV-, Flu-, and CMV-specific CD8^+^ T cells from donors I and J after two weeks of culture. Cells were gated on live, CD3^+^CD8^+^dextramer^+^ cells. c, Low number of differentially expressed genes (padj>0.05) comparing CD94^+^CD62L^−^ and CD94^−^CD62L^+^ progeny of CMV-specific CD8^+^ T cells suggest less biased clonal identities in chronic infection.

